# NAP-CNB: Bioinformatic pipeline to predict MHC-I-restricted T cell epitopes in mice

**DOI:** 10.1101/2020.10.05.327015

**Authors:** Carlos Wert-Carvajal, Rubén Sánchez-García, José R Macías, Rebeca Sanz-Pamplona, Almudena Méndez Pérez, Ramon Alemany, Esteban Veiga, Carlos Óscar S. Sorzano, Arrate Muñoz-Barrutia

**Author notes:** Equal contributor (Carlos Wert-Carvajal, Rubén Sánchez-García).

## Abstract

Lack of a dedicated integrated pipeline for neoantigen discovery in mice hinders cancer immunotherapy research. Novel sequential approaches through recurrent neural networks can improve the accuracy of T-cell epitope immunogenicity predictions in mice, and a simplified variant selection process can reduce operational requirements. We have developed a web server tool (<u>NAP-CNB</u>) for a full and automatic pipeline based on recurrent neural networks, to predict putative neoantigens from tumoral RNA sequencing reads. The developed software can estimate H-2 peptide ligands, with an AUC of 0.95, directly from tumor samples. As a proof-of-concept, we used the B16 melanoma model to test the system’s predictive capabilities, and we report its putative neoantigens. NAP-CNB web server is freely available at http://biocomp.cnb.csic.es/NeoantigensApp/ with scripts and datasets accessible through the download section.

## Introduction

Cancer cells can accumulate many mutations that change protein sequences. It can lead to MHC-restricted T-cell epitopes [1]. Identifying the tumor-specific epitopes that elicit T cell cytotoxic responses represents a major challenge for cancer immunotherapy, particularly to design personalized therapies [1, 2]. Finding neoantigens in every cancer patient will be fundamental for the next generation of antitumor immunotherapies.

A plethora of neoantigen discovery pipelines has been described to enable the prediction of epitopes from genetic information. However, current pipelines are human-centered and, thus, are primarily designed for clinical usage [3, 4]. Among the preeminent research lines, genomic analysis adjustments [3, 5–8], and neoepitope ranking practices [5, 6, 8, 9] have been prioritized over immunogenicity prediction algorithms. Despite this, the latter remains a critical component of the overall workflow for which limited available options exist [10].

The absence of dedicated tools for the alternative *in vivo* mouse models hinders pre-clinical cancer immunotherapy research. Hence, laboratories have to produce or adapt to *ad-hoc* human pipelines. Solely Epi-Seq [11] and MuPeXI [9, 12] offer modified versions for the murine model. Both platforms follow the canonical prediction process, based on sequencing data to estimate the gene expression and the affinity with the T-cell receptor (TCR) affinity of the mutated peptide [10], which is a prerequisite to eliciting an immune response [1]. However, Epi-Seq is not tailored to neoantigen detection. It was conceived instead for the discovery of common tumor antigens. MuPeXI lacks genome preprocessing and variant calling in its analysis. Mainly, in both cases, the algorithms underpinning immunogenicity prediction rely on dense neural networks. In the case of Epi-Seq, it uses NetMHCPan [13], which is trained with 9meric samples from the major histocompatibility complex (MHC) of mice or H-2, while MuPeXI employs NetH2pan [14], which is trained with pooled sequences from a diverse set of alleles.

Supervised machine learning methods have facilitated the identification of neoepitopes. In particular, artificial neural networks have proven to be highly efficient [15]. Among them, recurrent neural networks (RNN) are better suited for sequential problems, as attested by their extensive usage in natural language processing systems [16]. As a case, long short-term memory (LSTM) units are, at present, used for protein prediction of function and interactions [17, 18].

Prediction models have relied on gene expression information from tumor samples to determine putative peptides for intervention [1]. However, current approaches depend on genetic information from DNA sequencing to determine mutations [5, 8]. This dependence hinders temporal performance and increases intervention costs, but whole-exome sequencing (WES) is justified for its improved selectivity [19]. Hence, a system may rely exclusively on RNA sequencing (RNA-Seq) to simultaneously identify mutations and gene expression levels [19]. If compensatory methods in immunogenicity prediction are present, a tool designed for pre-clinical use may only rely on mutational information from RNA-Seq for a cost-effective solution.

We developed an integrated pipeline optimized for a murine model that finds putative neoepitope via next-generation sequencing (NGS) tumor variant calling and ranks them using LSTMs. This novel platform is only based on RNA-Seq, and is automated for a given haplotype. As a proof-of-concept, we trained our system with the H-2K^b^ haplotype (MHC class I) to be tested for the commonly used B16 melanoma model in C57BL/6 mice but the tool is compatible with additional typings. The resource NAP-CNB is freely available as a web server at biocomp.cnb.csic.es/NeoantigensApp/.

## Methods

The proposed pipeline employs genome preprocessing tools, variant calling software, and customized neural network architecture to obtain putative neoantigens from RNA-Seq experiments. As an integrative tool, the workflow has been adapted into a web server for RNA-Seq file submissions, as represented in **Figure 1a**. A tumor RNA-Seq file should be inputted as “.fastq.gz” together with the MHC class I type and an email address to receive the final results in less than ten hours.

**Figure 1.**
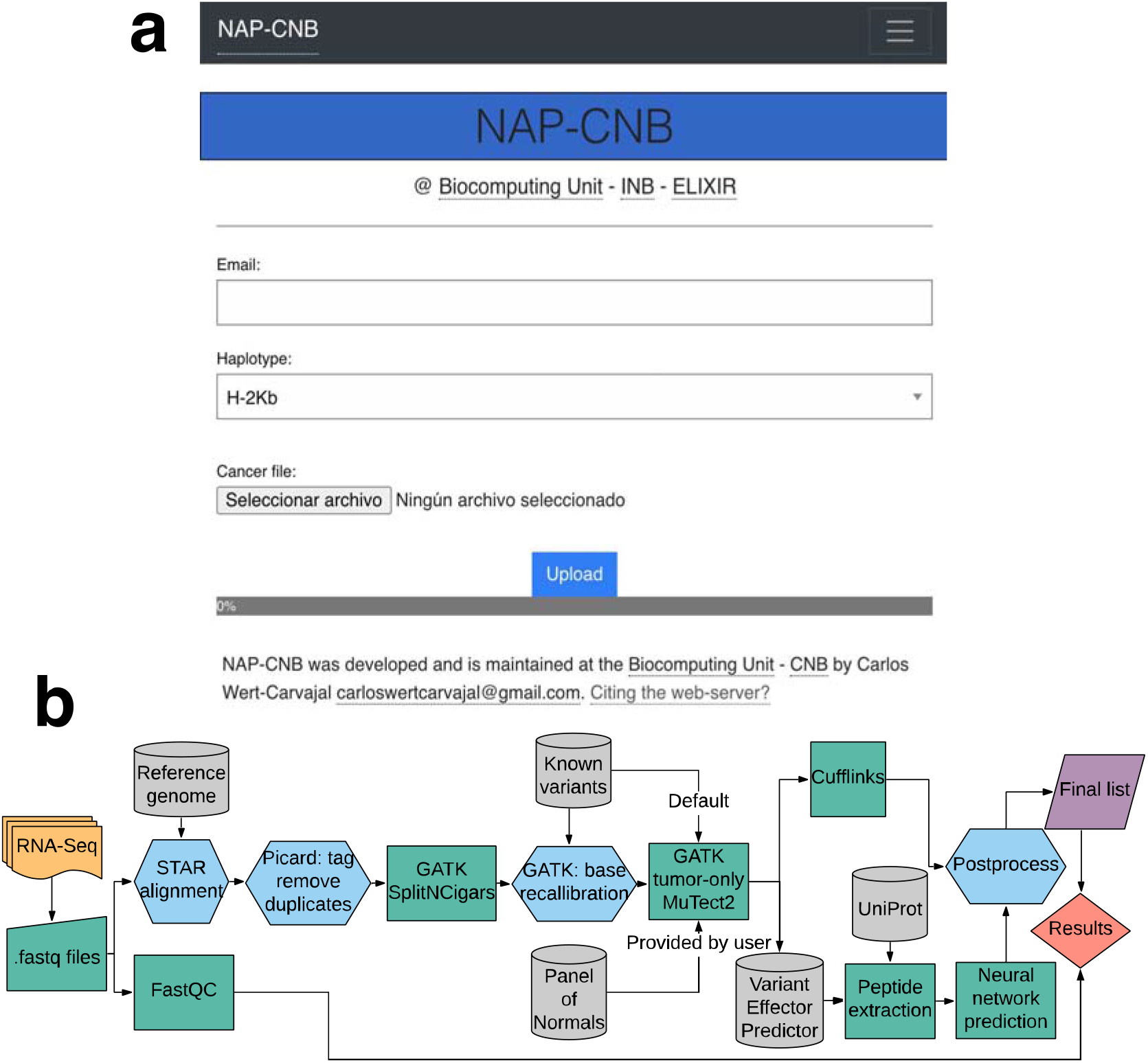
Workflow for the integrated pipeline. **(a)** The user interface of NAP-CNB (http://biocomp.cnb.csic.es/NeoantigensApp/) with the fields required for NGS analysis. Additionally, users may submit peptidic sequences for immugenicity prediction. Individual submissions are haplotype-specific, and results are sent to an email address. **(b)** Workflow for the integrated pipeline. Firstly, the sample is preprocessed before variant calling. Quality control through FastQC and STAR alignment with the reference genome is followed with protocols from Best Practices of GATK. Known variants are introduced through known polymorphisms or a panel-of-normals if requested, andsufficient non-tumor RNA-Seq reads are provided. MuTect2 is used for variant calling, and plausible single nucleotide variant (SNV) mutations translated into peptidic sequences for prediction with the RNN model. Gene expression is quantified through Cuffquant in Cufflinks.

### Variant calling: from RNA-Seq to mutant peptides

The somatic mutations suitable for neoantigen prediction are obtained from the gene expression of tumor tissue (RNA-Seq). NGS technologies that produce a FASTQ file are required for this protocol.

First, a quality assessment report is produced using FastQC (v0.11.8) [20] for user evaluation. In terms of preprocessing, the RNA-Seq file is realigned with a reference genome for further processing with STAR (v2.6.0a) [21]. The resulting BAM file is processed with Picard (v2.19.2) [22] for further refinements such as annotation and duplicate marking. Subsequently, Genome Analysis Toolkit (GATK, v4.1.2.0) [23] is used for exon segmentation, through the “SplitNCigarsReads” protocol, and base recalibration following Best Practices guidelines [24]. As indicated in **Figure 1**, this part serves as a preprocessing of the RNA-Seq reads *per se* before variant calling.

The MuTect2 variant caller [25] from the GATK package is used in its tumor-only mode, as shown in **Figure 1**, which is computationally less expensive but provides a higher number of false positives [26]. Even if designed primarily for DNA-Seq reads, MuTect2 has shown to be efficient in calling mutations from RNA-Seq [27]. By default, tumoral RNA-Seq is matched with databases of single nucleotide polymorphisms (dbSNP), although it can be used with a panel-of-normals (PoN) by construction. Following depth coverage filtering, the variants are submitted to Variant Effector Predictor (VEP) from Ensembl (v100.0) [28] for annotation and extraction of mutant peptide sequences identified as missense variants. Finally, a script matches the resulting UniParc reference from VEP to extracted UniProt proteins for protein-level prediction [29].

Additionally, Cufflinks (v2.2.1) [30] is used for mRNA abundance estimation as measured by fragments per kilobase million (FPKM). As there is no range for optimal neoantigen expression, this metric is provided to the user for its examination, as shown in **Figure 1**.

### Dataset generation and preprocessing

Sequences of immunogenic peptides for algorithm development were obtained from the IEDB database [31] for the H-2K^b^ MHC-I haplotype of the mouse strain C57BL/6. Given the different binding assessment methodologies considered in IEDB, elements were binarized by their MHC class I classification as positive or negative, per IEDB standards. The dataset, by entries accession number, is available at NAP-CNB.

Peptides deemed as antigenic were processed to extract their binding sites. Positive epitopes from IEDB were aligned with its protein source through the Smith-Waterman algorithm [32] to obtain the remaining sequence as negative samples (Suppl. Fig. 1). Additionally, epitope regions were extended through the original sequence to have a regular size (Suppl. Fig. 1). In contrast with previous methods, a given prevalence (i.e., the fraction of the minority class) was not imposed on the dataset. In total, 4,828 peptide entries were processed into 251,049 sequences with 6,714 positive entries and 244,225 negatives. A 10% split was used for test set generation. Concerning blind test data, IEDB datasets 1034799 and 1035276 were processed through the previous procedure and by the method described by [14].

Further postprocessing was implemented with an optional majority vote algorithm that considered mutations to the most similar amino acid, given by the BLOSUM62 matrix [33], for each position. In other terms, a sequence modified its classification if there was a consensus among its most akin peptides.

### Neural network training

The neural networks were implemented through Keras (v2.2.4) [34] and TensorFlow (v1.11.0) [35]. A scalable routine was used for architecture optimization through simplified datasets (Suppl. Fig. 1) until one competent was obtained. Moreover, training was done with “on-batch” class balancing and data augmentation. The latter increased the number of positives sequences through random substitution of a given number of amino acids with similar ones from the BLOSUM62 matrix [33], with a given tolerance (Suppl. Fig. 3). The training was performed through 5-fold cross-validation, for hyperparameters tuning and optimization of balancing and augmentation, generating a total of 80 models for the actual dataset.

The initial toy model was used for embedding selection and model tuning of neural architectures (Suppl. Table 1A-B), which was maintained in the type and depth of layers in later configurations. While an intermediate dataset (Suppl. Fig. 1C) was introduced for data balancing and augmentation. The final model was produced with the complete dataset and cross-validation of the number of internal LSTM units at each layer, the number of on-batch sequence augmentations, and its tolerance, and the on-batch class balancing.

In the final architecture, 12mer peptide sequences are introduced with a one-hot encoding representation to three consecutive bidirectional LSTM layers, followed by three layers of dense neurons with two intermediate dropouts units. The output layer consists of a dense neuron, with a soft-max activation, which yields the affinity estimation probability. The overall network is represented in **Figure 2**.

**Figure 2.**
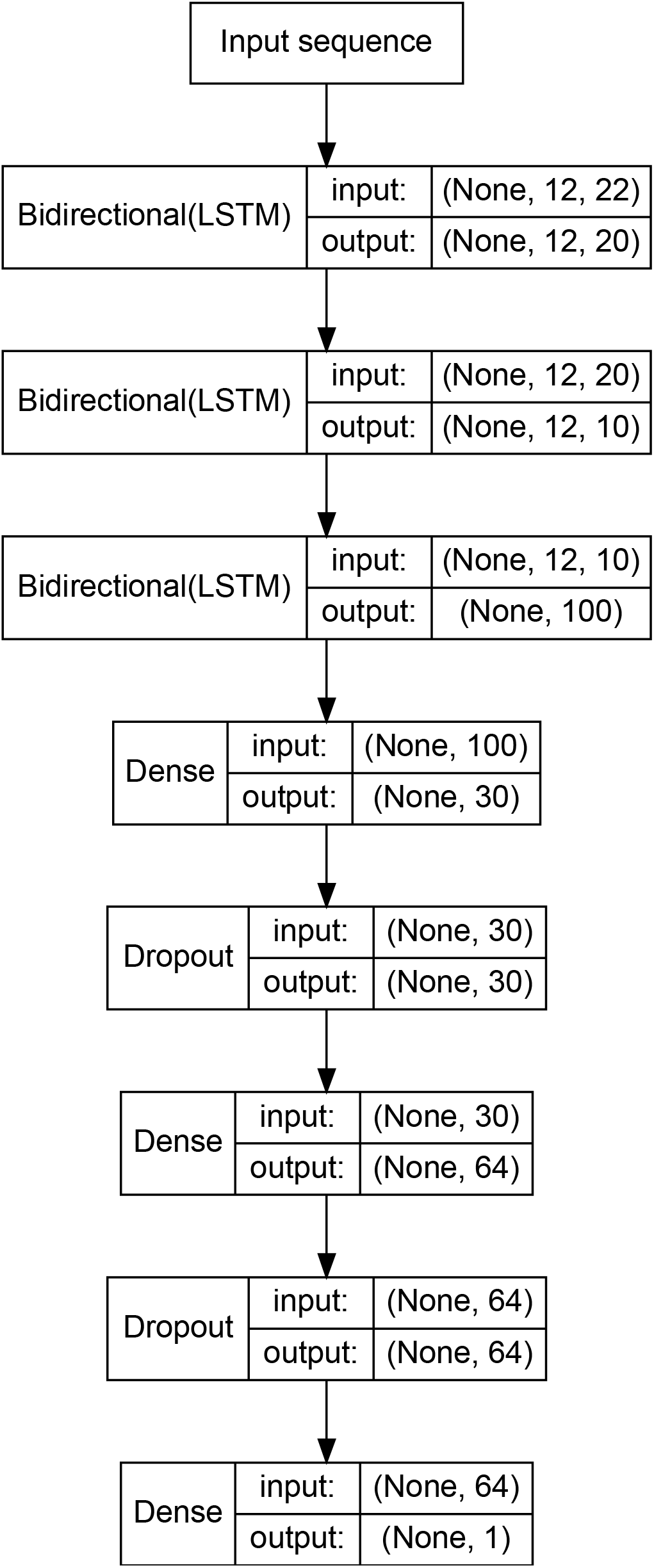
Neural network model of the affinity prediction for H-2K^b^. The input sequence corresponds to a one-hot encoding of a 12mer peptide sequence extracted from the preprocessing workflow. The number of LSTM units corresponds to the input sequence’s overall length across the three consecutive layers. Following the RNN, two hidden dense units, with alternating dropouts, serve to process an affinity probability.

### Sequencing raw data

An *in vitro* B16 melanoma cell line with an H-2K^b^ haplotype was processed for RNA extraction and sequenced through an NGS Illumina HiSeq2000. From the FastQC analysis, all evaluated parameters were satisfactory except from the presentation of four over-represented sequences corresponding to Illumina single end PCR primer and technical noise as TrueSeq adaptors. Trimming of these sequences was done before RNA-Seq processing. The resulting “fastq.gz” file was introduced for analysis in a local server.

## Results

### Cross-validation metrics

Initial architectures, based on LSTM and dense layers, showed performance improvements, in terms of the area under the curve for the receiver operator characteristic (AUC ROC), for higher depth models (Suppl. Table 1A). Despite this, these changes did not have an impact as significant as “on-batch” balancing and data augmentation. In particular, modifications of a “virtual” prevalence raised AUC ROC and F-1 values to 20% in test sets (Suppl. Table 1C) and decreased the degree of overfitting. All parameters were adjusted through grid search on the final model under a limited number of epochs (see Additional file 2 - Grid search parametrization). As observed in **Table 1,** the network’s final AUC reached 95%, albeit with an acceptable F1 score, due to the assumed low prevalence. The complete cross-validation results of each model are available at NAP-CNB.

**Table 1.** Binary classification metrics for the final 5-fold cross-validated algorithm. The reported mean statistics estimators correspond to AUC ROC, accuracy (ACC), precision or positive predictive value (PPV), and sensitivity and specificity with their harmonic average (F1). The prevalence of positive samples was around 1:40.

| AUC ROC | ACC | PPV | Sensitivity | Specificity | F1 |
| --- | --- | --- | --- | --- | --- |
| (±SD) | (±SD) | (±SD) | (±SD) | (±SD) | (±SD) |
| $0.95 \pm 0.04$ | $0.977 \pm 0.004$ | $0.6 \pm 0.1$ | $0.62 \pm 0.09$ | $0.988 \pm 0.004$ | $0.6 \pm 0.1$ |

For further evaluation, 10% of the original dataset was used as a test set of the selected parametrized system. In **Figure 3**, both the ROC and the precision-recall curve are shown. The latter reflects how the system fares against a high-class imbalance. In terms of metrics, the ROC AUC for the test sample was 86.5% with 97.2% accuracy. Notwithstanding, the proposed ensemble method for postprocessing could increase precision by 7.6%.

**Figure 3.**
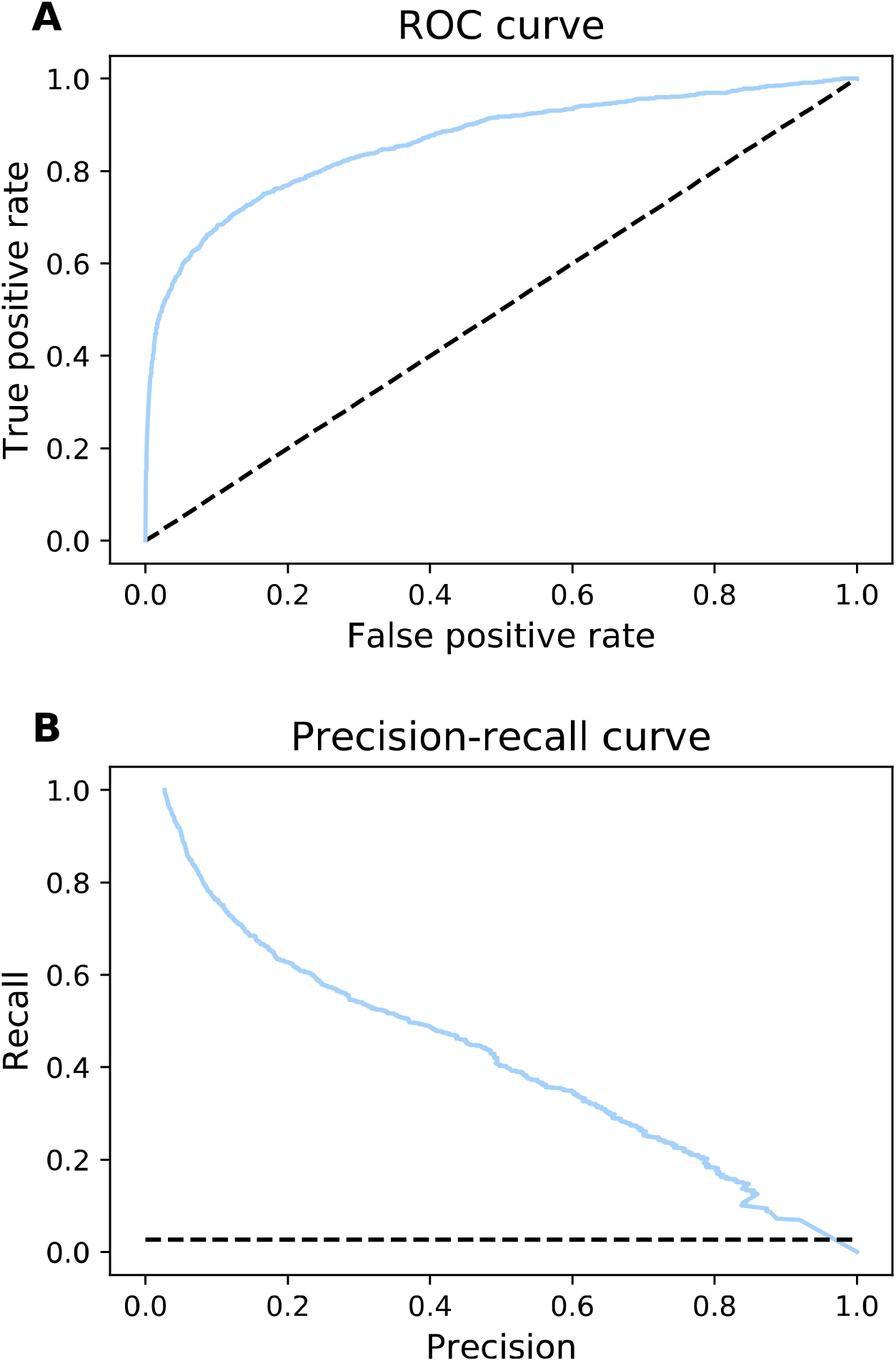
ROC and precision-recall curves for the final model. **A)** ROC curve for 10% test partition with an AUC of 86.5%, the dashed line shows chance level. **B)** Precision-recall curve with the prevalence of around 3% shown as chance. The precision-recall AUC is 41.97%, whereas a random guess corresponds to an AUC of 2.64% for the same data imbalance.

Throughout cross-validated models, window sizes of 8, 10, and 12 amino acids were tested for predictive performance. Sequences of 12 amino acids produced more accurate models, as observed in **Figure 4**. This result may indicate that antigenic determinants are not sufficient for peptide classification and distal amino acids carry additional predictive information. The distribution of sequences classified as positive and a sensitivity analysis from random classifications showed similar results (Suppl. Fig. 4). In the typing H-2K^d^, the best performance corresponded to 8-mers, which is the window used for mutant peptides (Suppl. Material H2-Kd). Cross-prediction between both typings was suboptimal (Suppl. Material. H2-Kd), which may indicate that neural networks should be configured for each typing. The cross-validation metrics of the additional haplotypes are also available (Suppl. Material. H-2 Metrics), showing both enhancements and reductions in efficacy.

**Figure 4.**
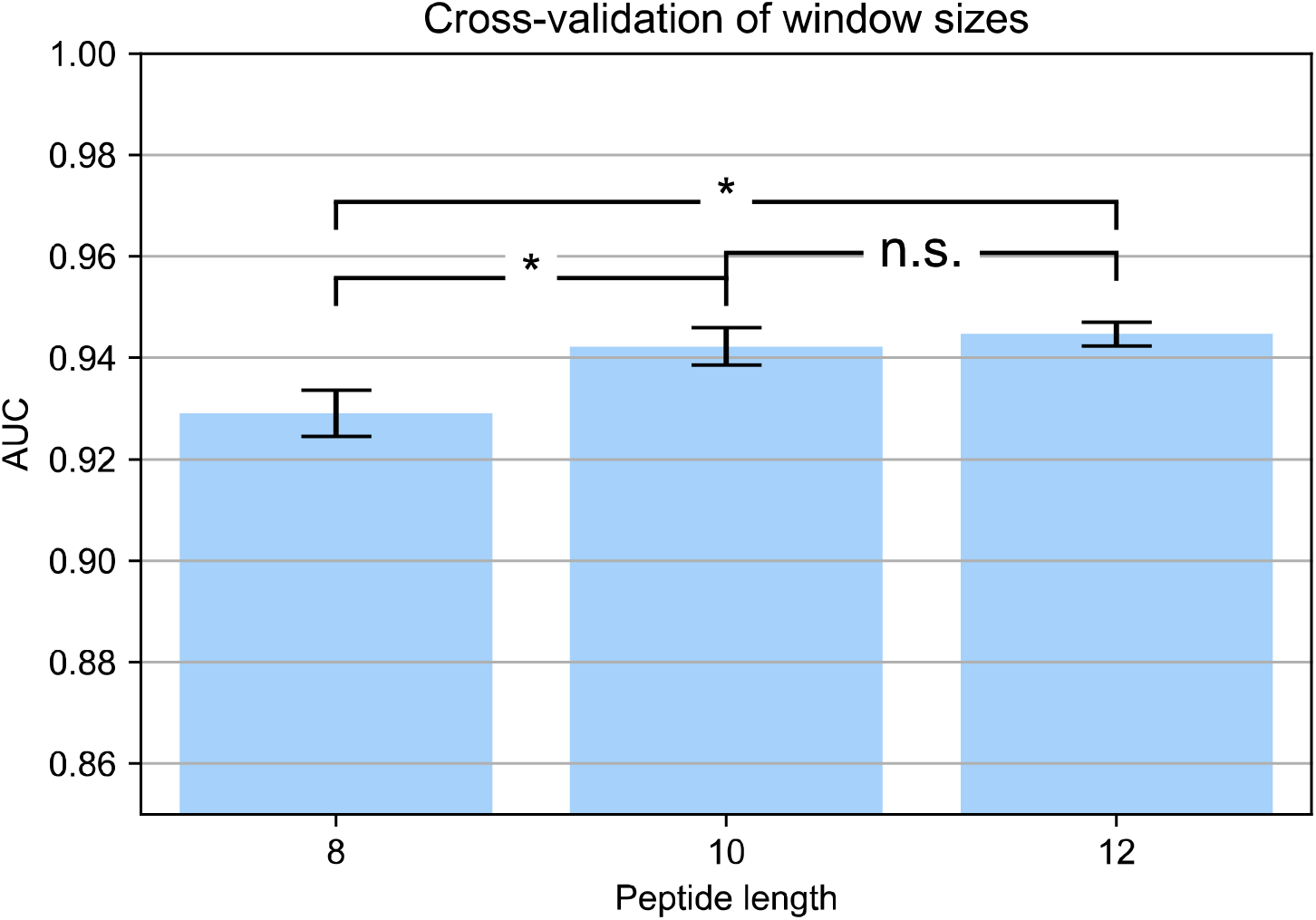
Cross-validation of peptide window sizes. The area under the curve of the receiver operating characteristic curve using 8mers, 9mers, and 12mers obtained through 5-fold cross-validation in different conditions. The windows are obtained from the mutated peptide sequence centered at the location of the SNV. Significant differences between means (p<0.05) are shown.

### Benchmarking

Compared against NetH2pan [14], which is the benchmark used for MHC class I affinity prediction in mice, the reported cross-validated AUC ROC of 95% is 3% higher for the H-2K^b^ typing and a similar performance in PPV. To confirm a better performance on a dataset, blind testing was implemented from two new H-2K^b^ datasets from IEDB (1034799 and 1035276). Negatives were generated following the protocol mentioned above, disregarding positive sequences that do not have a protein accession or cannot be reframed into 12-mers, and by generating random sequences with an assumed prevalence as described in [14]. Given that NetH2Pan considers different epitope lengths and substitutions, binarization was done by considering whether binds were predicted overall for a 12mer sequence. In all binary metrics, the LSTM network achieved improved results (Suppl. Fig 5 and 6). The reported accuracies were between 96% and 98%, with up to 3-fold increases in precision.

Notably, in all cases, positives were better detected than in NetH2pan. All this irrespective of the method used to produce negative sequences. On the whole, our approach detected 259 and NetH2pan 86 of a total of 438 antigens across both datasets. Moreover, an ensemble method joining predictive positives from both methods improved detection to 277 with random negatives and 254 with negative sampling.

### Use case

As a result of MuTect2 calling, 4,566 variants were identified. From those, 1,085 missense transcripts were obtained from VEP corresponding to 345 genes. These were matched against the results from Cufflinks and submitted for prediction. In the end, our proposed software generated a ranking of putative neoantigens.

The thirty-five top-scoring putative neoepitopes are shown in **Table 2**. The predictions were matched with the original B16 results from Castle *et al.* [36] (Suppl. Table 2). Additionally, we compared the rank given by our proposed algorithm’s softmax score with the relative classification of the 12mer sequence in NetH2pan, obtained by averaging the scores across all of its possible epitope lengths and mutations [14]. **Table 2**, thus, establishes an order of preference for both methods. Due to sample size limitations, the haplotype H-2D^b^ of the C57BL/6 model is not analyzed but should also be included in a naïve study.

**Table 2.**
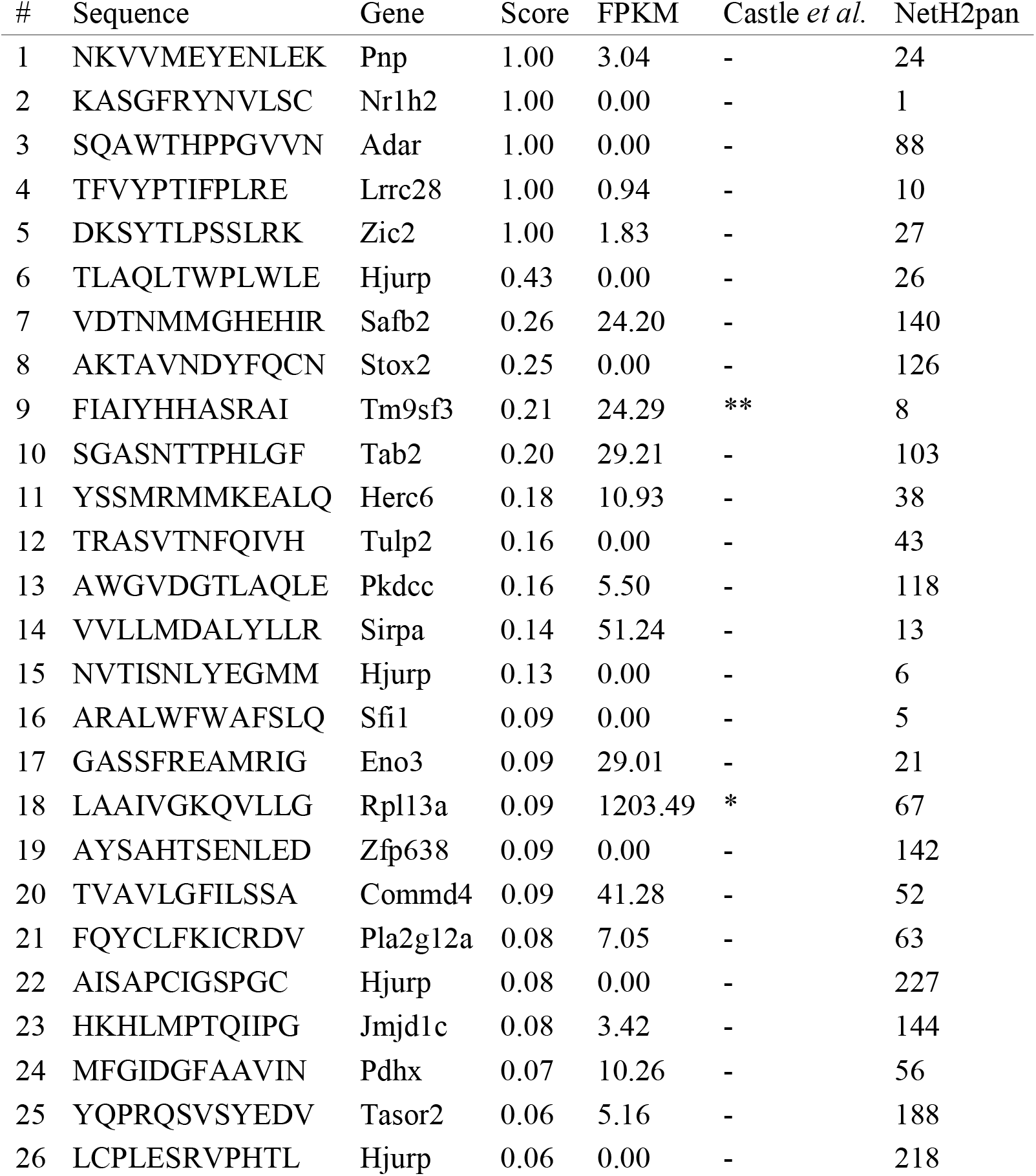
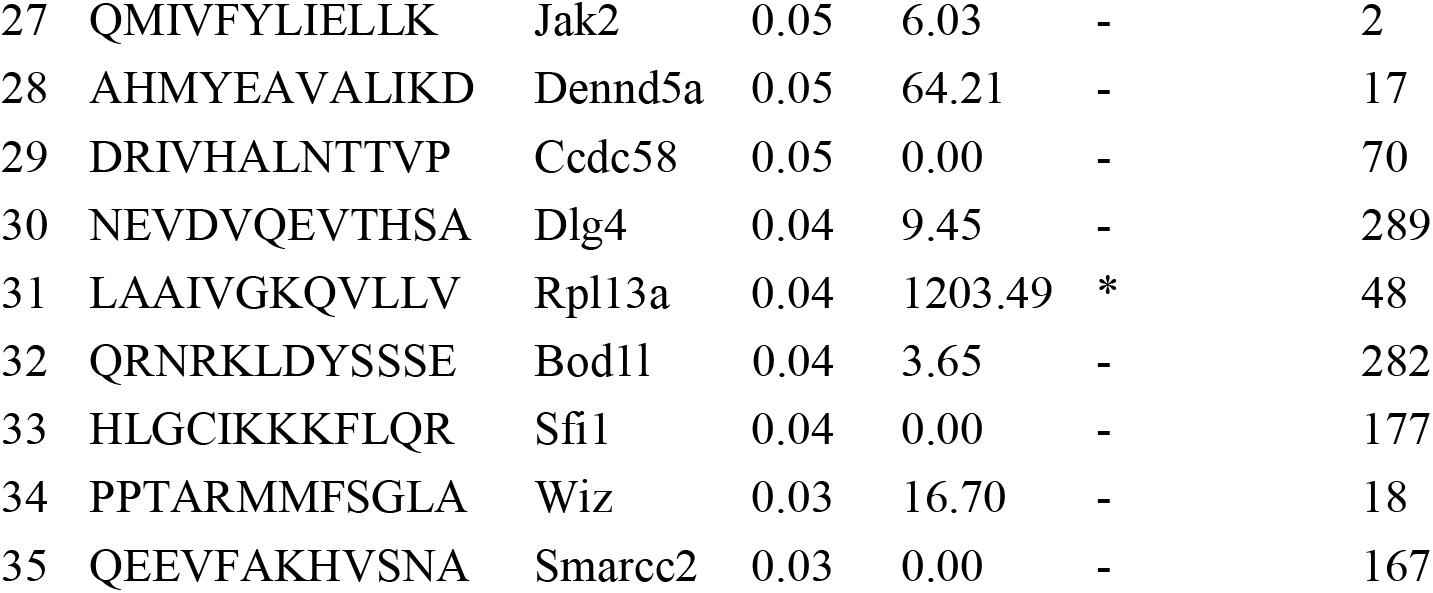
Putative neoantigens, shown by sequence and gene symbol, for the B16 melanoma model, restricted to H-2K^b^, ranked by scores. The gene expression is quantified as fragments per kilobase million. Neoantigens examined in Castle et al. [36] are classified by selection for validation (*) and reactivity (**). The classification of the average score for a complete 12mer sequence given by NetH2pan is also presented.

In terms of overall performance, the entire pipeline has an execution time of around eleven hours in a local server using two CPU cores. This duration corresponds to steps between preprocessing of the RNA-Seq and quality analysis to immunogenicity prediction. The levels of abundance may be suboptimal in some cases. Still, its reporting may guide the user in selecting a candidate.

## Discussion

The proposed pipeline provides an integrated software solution for mouse neoantigen MHC class I discovery from RNA-Seq data. The workflow is based on a streamlined process adapted to the resource-efficient and accessibility requirements of pre-clinical research. Notably, we report an immunogenicity estimation model that successfully improves previously reported performance. The B16 case study also shows a good number of putative neoantigens that are coherent with literature estimates [36]. A functional validation measuring T-cell immune responses by ELISPOT or intracellular IFN-gamma staining in mice responding to B16 tumors would be required to validate the prediction results.

In terms of the actual prediction algorithm, the RNN-based approach presents an AUC of 95% in cross-validation. Compared with the current NetH2pan benchmark model [14], it represents an enhancement in terms of accuracy and precision for the H-2K^b^ haplotype in both cross-validation and blind testing metrics, with a 3-fold increase of precision in the latter. However, this varies depending on the haplotype used, with H-2K^d^, for instance, lacking such improvements for a blind set. Thus, these results may reinforce sequential models’ usefulness as an efficient solution to antigen binding prediction against more conventional neural network approaches. Future lines of research may include more recent sequential model innovations. Novel types of sequential architectures in transformers and RNNs, such as BERT [37] and GORU [38], could serve as enhancers of overall performance. Also, subsequent work in epitope size should aim to reconcile flexibility, which is compatible with an RNN-based framework, with the generation of empirical negative samples. The web server restricts the haplotype utilized for prediction. Even if cross-prediction between haplotypes K^b^ and K^d^ suggests type-specific modeling is an optimal solution, a pan-specific system is part of future directions.

Concerning data processing, the use of negative empirical sequences and data augmentation should also be considered to improve affinity estimation. Strategies could include generative models such as Gaussian mixtures or adversarial networks (GAN) [39]. Nonetheless, one of the problems posed by the dataset is its reliance on a binarized predictor which hampers the biological meaning of the results. Another problem is the prevalence dependency of precision and recall. Further work should be done to identify an optimal strategy. Finally, our method is characterized by the employment of window sizes that are above the normative length of an epitope to optimize performance, which may imply that reported antigenic determinants are not sufficient information for prediction. From a biological perspective, this is equivalent to considering the overall conformation of a protein as relevant in the binding process.

The variant calling process poses further challenges. Our approach has prioritized a procedure that functions solely on RNA-Seq data with a conservative selection of mutations, particularly missense SNV. This neglects a high percentage of variants that produce neoantigens [40] and increases the mutational uncertainty by not including genomic data from DNA-Seq [19]. Advances should proceed in this direction, albeit prioritizing an exclusive RNA-Seq utilization to retain the tool’s cost-effectiveness, which is essential for our open web service to remain reachable.

## Supporting information

Grid search parametrization

## Additional Files

**Additional file 1 — Supplementary material**

**Additional file 2 — Grid search parametrization**

