## Supplementary material for "NAP-CNB: Bioinformatic pipeline to predict MHC-I-restricted T cell epitopes in mice"

### Supplementary Figure 1

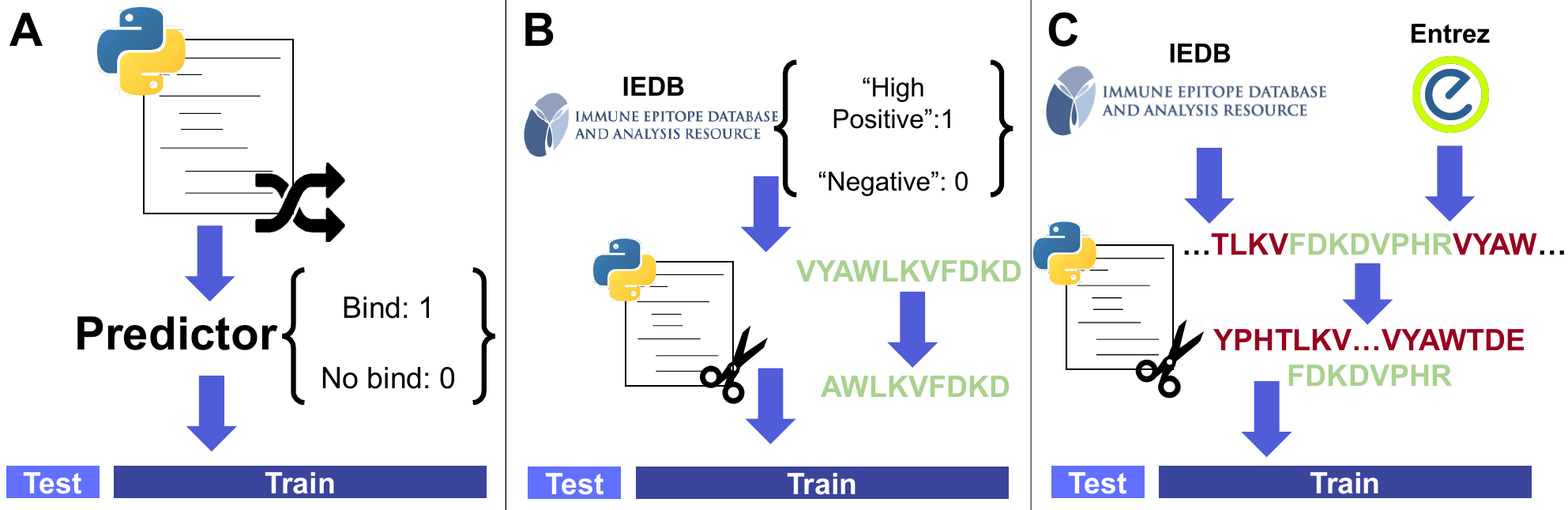

A) Initial architecture configurations used random peptides with the binarized classification from NetH2pan as a toy model. B) For data augmentation and balancing trials, the dataset consisted of epitopes categorized by IEDB as “high positive” and “negative” entries with an equal window. C) In the final model, epitopes were extracted as 12-mers from the original protein, with the rest of the peptide extracted as negatives.

### Supplementary Figure 2

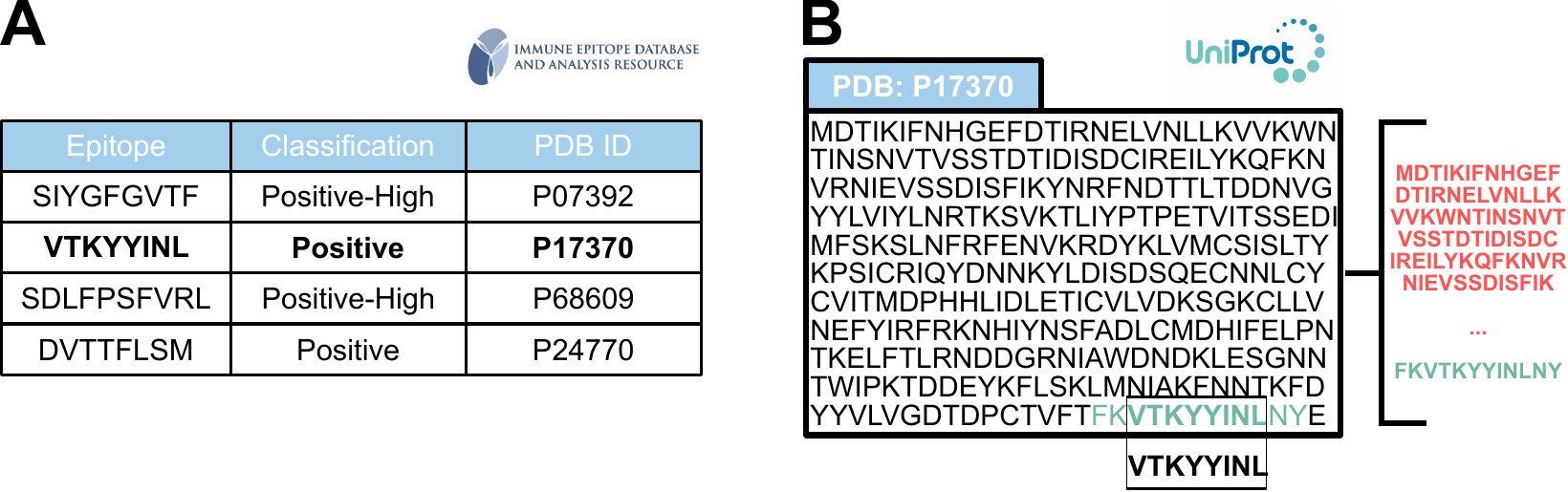

Data from IEDB (A) is aligned through the Smith-Waterman algorithm with the PDB entry from UniProt to obtain an extended sequence and get negatives for training with the remaining sequence. (B) These sequences are then balanced on-batch for different prevalence levels.

### Supplementary Figure 3

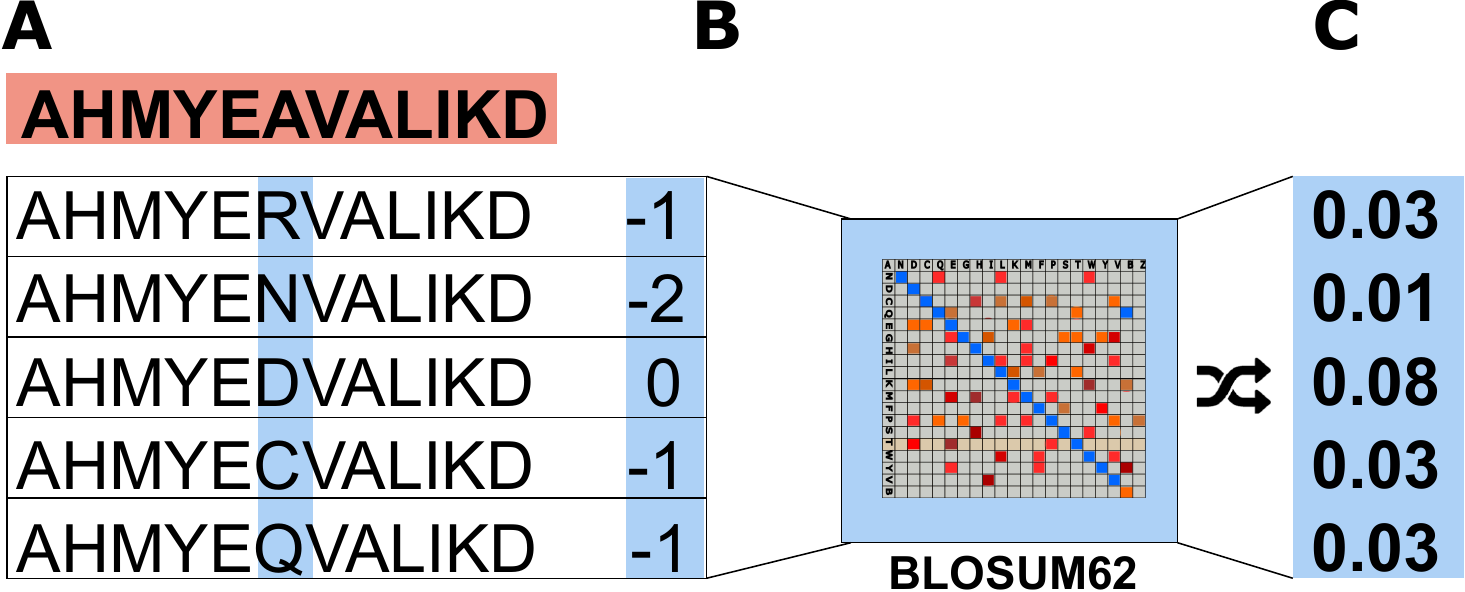

For a sequence (A), data is augmented through mutations at a random location. The new amino acid's similarity score, extracted from the BLOSUM62 matrix (B), normalized using a softmax function (C), is higher than a given tolerance. The number of amino acids to be mutated and the proportion of sequences per batch to be augmented serve as additional optimization parameters.

### Supplementary Table 1

**A)**

| **#LSTM** | **Output** | **AUC** | **Precision** | **ACC** | **Sensitivity** | **Specificity** | **F-1** |
| --- | --- | --- | --- | --- | --- | --- | --- |
| 1 | 10 | 0.978 | 0.994 | 0.972 | 0.954 | 0.993 | 0.999 |
| 2 | 5 | 0.999 | 1.000 | 0.995 | 0.997 | 0.993 | 0.996 |
| 3 | 5 | 1.000 | 0.994 | 0.984 | 0.976 | 0.993 | 0.985 |

Tests on different depths in long short-term memory units for a batch size of 10 and 5 epochs. We considered the best results in terms of output size in the range of 5,10, or 20. The training was performed on the toy model (Supplementary Figure 1A). Metrics correspond to a 10% test set.

**B)**

| **Embedding** | **Dim** | **AUC** | **Precision** | **ACC** | **Sensitivity** | **Specificity** | **F-1** |
| --- | --- | --- | --- | --- | --- | --- | --- |
| One-hot | 22 | 1.000 | 0.976 | 0.987 | 1.000 | 0.972 | 0.988 |
| Kidera *et al.* [1] | 10 | 0.999 | 0.991 | 0.982 | 0.976 | 0.990 | 0.983 |
| Liu *et al.* [2] | 11 | 0.998 | 0.991 | 0.972 | 0.957 | 0.990 | 0.974 |
| Atchley *et al.* [3] | 5 | 0.999 | 0.973 | 0.982 | 0.994 | 0.969 | 0.984 |

Performance metrics of embeddings extracted from the literature for amino acid representation. The toy model (Supplementary Figure 1A) was not optimized and parametrized for each case, using the fittest for the one-hot encoding for comparison. Metrics correspond to a 10% test set.

**C)**

| **On batch additions** | **Tolerance** | **AUC** | **Precision** | **Sensitivity** | **Specificity** | **F-1** |
| --- | --- | --- | --- | --- | --- | --- |
| 0 | - | 0.937 | 0.763 | 0.766 | 0.943 | 0.765 |
| 1 | 0.00 | 0.957 | 0.767 | 0.838 | 0.941 | 0.801 |
|  | 0.05 | 0.947 | 0.774 | 0.689 | 0.952 | 0.729 |
|  | 0.10 | 0.934 | 0.726 | 0.796 | 0.926 | 0.759 |
| 2 | 0.00 | 0.943 | 0.737 | 0.818 | 0.931 | 0.775 |
|  | 0.05 | 0.931 | 0.701 | 0.718 | 0.927 | 0.709 |
|  | 0.10 | 0.919 | 0.668 | 0.711 | 0.901 | 0.689 |
| 5 | 0.00 | 0.948 | 0.754 | 0.826 | 0.939 | 0.788 |
|  | 0.05 | 0.934 | 0.756 | 0.593 | 0.955 | 0.665 |
|  | 0.10 | 0.933 | 0.713 | 0.713 | 0.931 | 0.713 |

Measures from the data augmentation tests for different numbers of new peptide entries per batch at different tolerance thresholds. A batch size of 20 entries and 20 epochs was used. Tolerance denotes the maximum normalized BLOSUM62 similarity for augmentation (Supplementary Figure 3). Thus, only mutations higher than the tolerance score were allowed. Augmentation changes were tested for method validity on the intermediate high-confidence model (Supplementary Figure 1B) with the mean of 5-fold cross-validation shown for each metric. See Additional file 2- Grid search parametrization

### Supplementary Figure 4

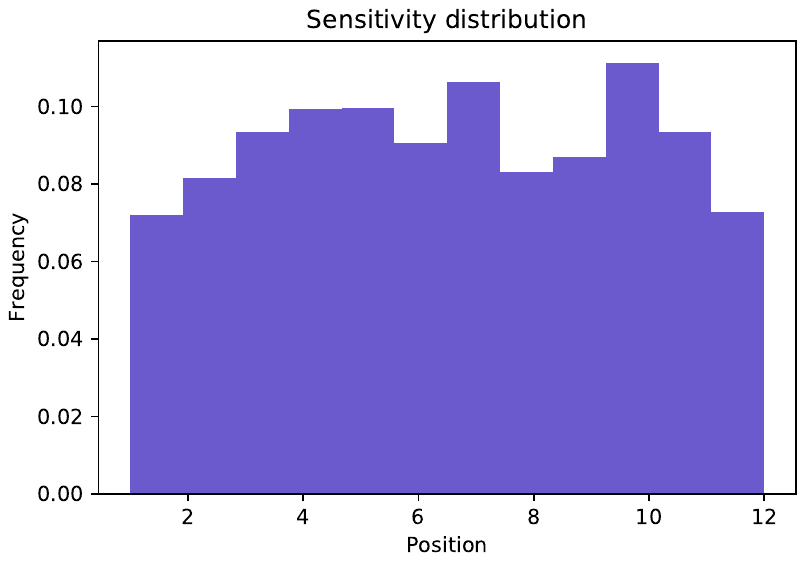

To characterize the susceptibility of each location to change the outcome, we generated 60,000 natural random peptides and produced random substitutions for each position. As a result, 5,843 sequences representing a 9.74% of the entire series, were prone to modify their prediction through a single amino acid variation. Of those, 83.04% altered their label from negative to positive. The resulting histogram failed to pass a two-sided Kolmogorov-Smirnov test for a uniform distribution (D = 0.097911, p< 2.2e-16), which implies sensitivity is not evenly distributed.

### Supplementary Figure 5

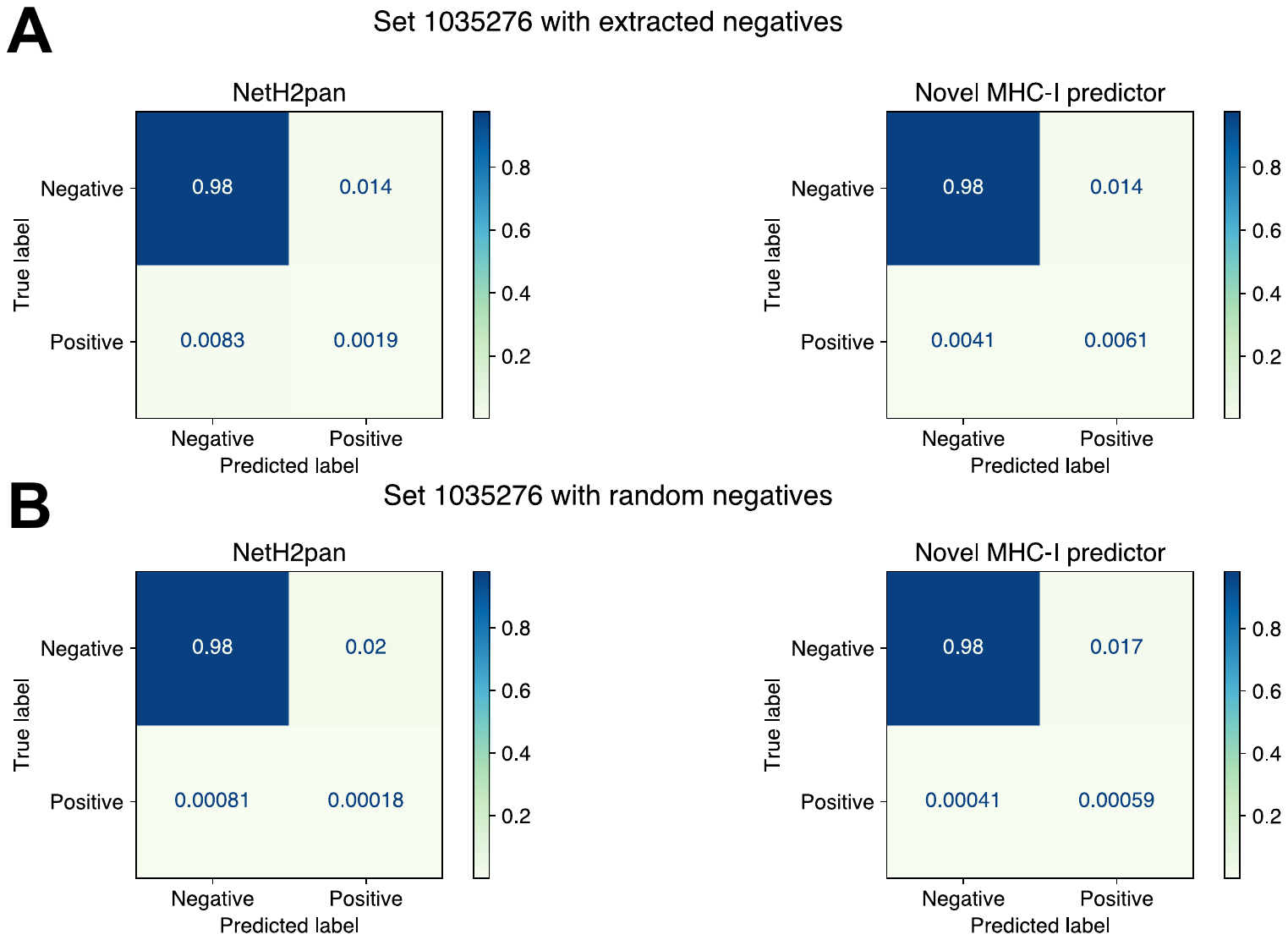

A) Normalized confusion matrix for negatives extracted entries in the 1035276 IEDB dataset. PPV 29.70% (NAP-CNB) and 12.01% (NetH2pan), ACC 98.15% (NAP-CNB) and 97.79% (NetH2pan). B) Negatives obtained from random sequences introduce 999 natural random peptides per positive sequence. PPV 3.33% (NAP-CNB) and 0.89% (NetH2pan), ACC 98.24% (NAP-CNB) and 97.89% (NetH2pan).

### Supplementary Figure 6

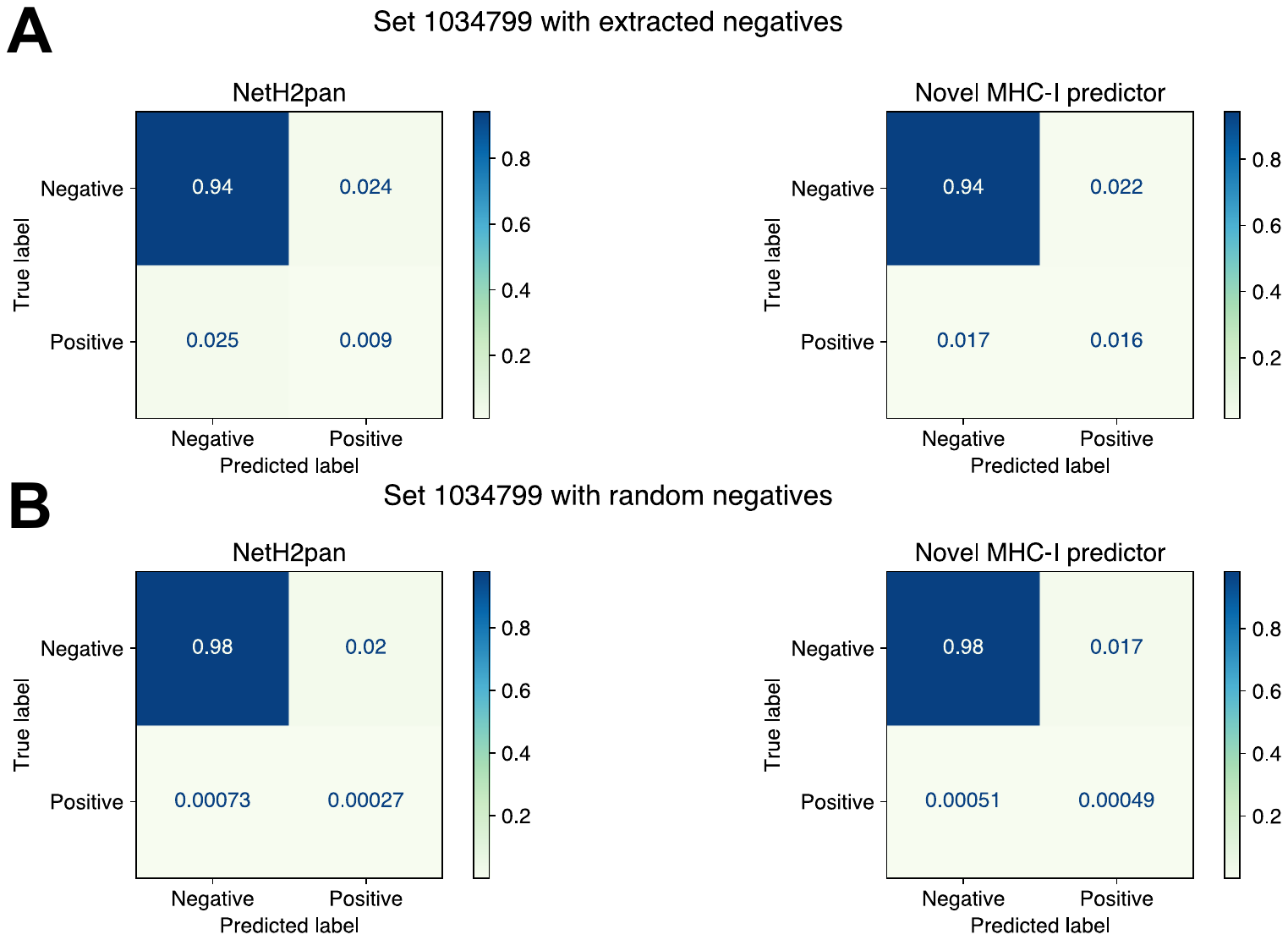

A) Normalized confusion matrix for negatives extracted entries in the 1034799 IEDB dataset. PPV 42.31% (NAP-CNB) and 27.27% (NetH2pan), ACC 96.03% (NAP-CNB) and 95.13% (NetH2pan). B) Negatives obtained from random sequences introduce 999 natural random peptides per positive sequence. PPV 2.81% (NAP-CNB) and 1.30% (NetH2pan), ACC 98.26% (NAP-CNB) and 97.90% (NetH2pan).

### Supplementary Table 2

| **#** | **Sequence** | **Gene** | **Score** | **FPKM** | **NetH2pan** |
| --- | --- | --- | --- | --- | --- |
| 9 | FIAIYHHASRAI | Tm9sf3 | 0.21 | 24.29 | 8 |
| 55 | WYTGEAMDEMEF | Tubb3 | 0.01 | 87.0 | 237 |
| 64 | TQLKKPFLVNNK | Ppp1r7 | 0.01 | 8.03 | 128 |
| 84 | FVDWENVSPELN | Kif18b | 0.01 | 3.32 | 143 |
| 158 | TTTKKARVSTPK | Dag1 | 0.0 | 0.0 | 259 |
| 166 | QAFIDVMSRETT | Actn4 | 0.0 | 87.84 | 150 |
| 248 | HLNNDVWQIFEN | Plod2 | 0.0 | 0.0 | 204 |
| 253 | GQQLVIQLLHTC | Tnpo3 | 0.0 | 39.04 | 75 |
| 283 | LVLHVVSAAQAE | Sema3b | 0.0 | 31.53 | 132 |

Complete validated immunogenic mutations from the original paper by Castle *et al.* [4] with the ranking of the mean score given by NetH2pan and the proposed LSTM-based algorithm. Also shown are the fragments per kilobase million (FPKM) of the gene expression.

**Supplementary Material for H-2K^d^**

The total dataset for a window of 12 peptides contained 1,531 positives and 63,686 sequences in total. Using this dataset for prediction with the ANN built for H2Kb, we obtained the binary metrics:

| ACC | PPV | Sensitivity | Specificity | F-1 |
| --- | --- | --- | --- | --- |
| 0.964 | 0.114 | 0.076 | 0.986 | 0.091 |

It corresponds to a 12mer peptide window with positives obtained from T-cell and MHC ligand assays from IEDB that had a protein entry with the epitope.

Thus, to improve the performance for low positive detections, we trained a different NN for this specific haplotype. Under different network configurations, 8mers systematically outperformed 10mers and 12mers in AUC ROC and PPV in 5-fold cross-validation. The parameters employed for H-2K^b^ did not produce a good performance after re-training; thus, we use a 5-fold cross-validation routine for optimization.

The dataset used for the 8mer sequences contained 1895 positive sequences and 93281 negative ones.

The cross-validation metrics of the final model were:

| AUC ROC | ACC | PPV | Sensitivity | Specificity | F-1 |
| --- | --- | --- | --- | --- | --- |
| (±SD) | (±SD) | (±SD) | (±SD) | (±SD) | (±SD) |
| 0.96±0.05 | 0.982±0.008 | 0.5±0.2 | 0.7±0.2 | 0.987±0.008 | 0.6±0.2 |

For the test set, the binary metrics are:

| ACC | PPV | Sensitivity | Specificity | F-1 |
| --- | --- | --- | --- | --- |
| 0.974 | 0.397 | 0.485 | 0.984 | 0.436 |

Due to its entry date and identification of the original peptides, we identified set 1036855 for blind testing with 9 positives. We generated 92 negatives from our method and 8991 from random negatives.

For this set, our method had less optimal results in overall positives identification. In comparison, NetH2Pan identified one epitope, whereas our approach did not predict any.

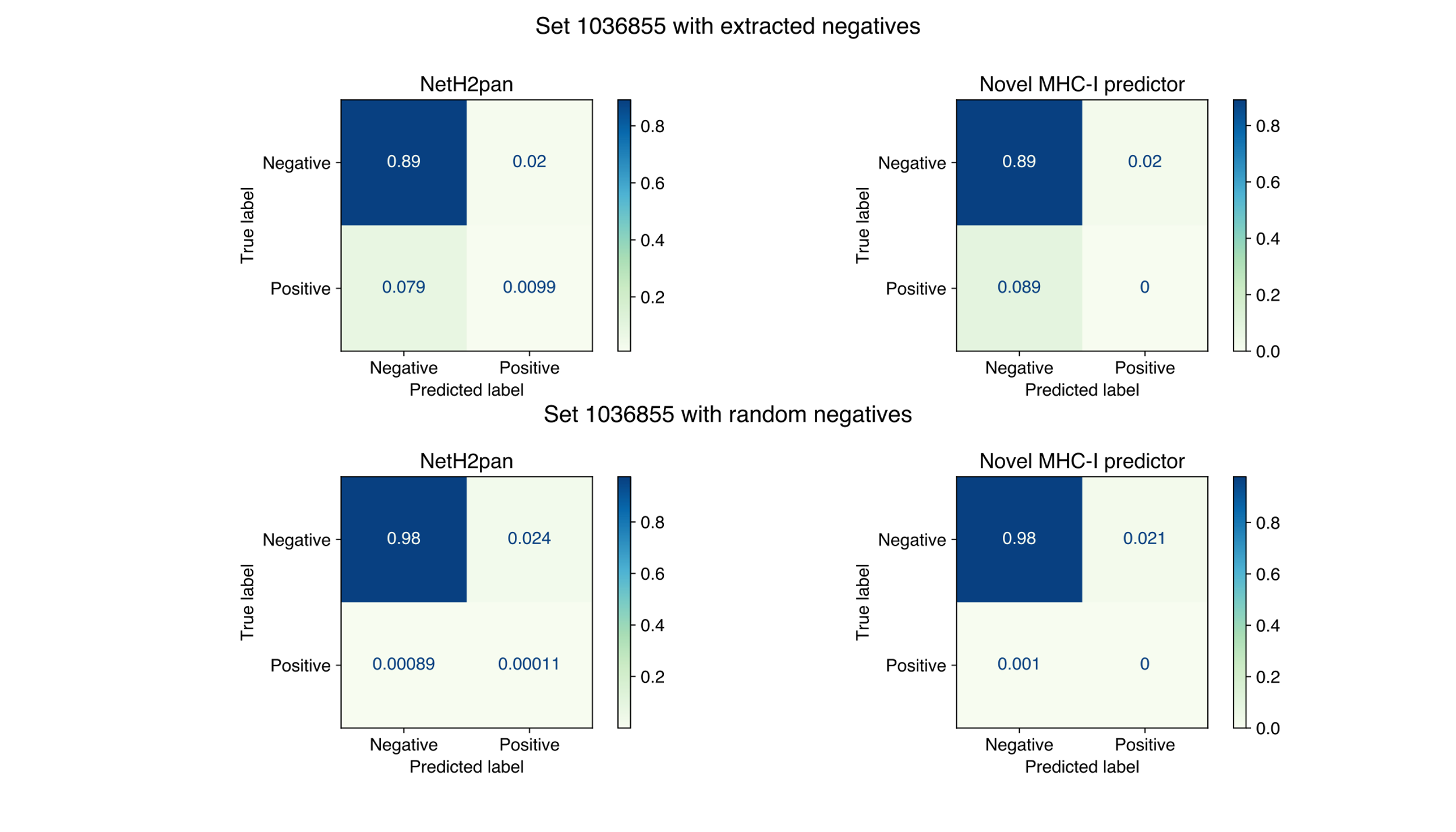

**Supplementary Material for other H-2 types**

The reported AUC ROC of the other H-2 haplotypes are:

| Haplotype | AUC ROC (±SD) |
| --- | --- |
| H-2Db | 0.7±0.1 |
| H-2Dd | 0.9±0.1 |
| H-2Dq | 0.8±0.1 |
| H-2Kk | 0.96±0.06 |
| H-2Kq | 0.9±0.2 |
| H-2Ld | 0.9±0.1 |
| H-2Lq | 0.7±0.2 |

Other binary metrics and dataset characteristics can be downloaded from the webserver.
